# Molecular and developmental signatures of genital size macro-evolution in bugs

**DOI:** 10.1101/2022.05.23.493079

**Authors:** Bruno C. Genevcius, Denis C. Callandrielo, Tatiana T. Torres

## Abstract

With the advance of the *evo-devo* research program, our understanding of the genes and pathways that determine the architecture of novel traits has experienced drastic growth. Nevertheless, single-species approaches are insufficient to understand the processes by which evolution shapes morphological changes after their emergence. As such, we still have an elusive knowledge of how these genetic-developmental architectures evolve themselves for most of the structures, as well as how their evolution is mirrored in the phenotypic change across large time scales. Here, we tackle this gap by reconstructing the evolution of male genital size, one of the most complex traits in insects, together with its underlying genetic architecture. Using the order Hemiptera as a model, which spans over 350 million years of evolution, we estimate the correlation between genital size and three features: development rate, body size, and rates of DNA substitution in 68 genes previously associated with genital development. We demonstrate that genital size macro-evolution has been largely dependent on body size and weakly influenced by development rate and the phylogenetic history. Our results further revealed positive correlations between mutation rates and genital size for 18 genes. Interestingly, there is great diversity in the function of these genes, in the signaling pathways that they participate in, and in the specific genital parts that they control. These results suggest that fast genital size evolution has been enabled by molecular changes associated with diverse morphogenetic processes, such as cuticle composition, patterning of sensory apparatus, and organ growth itself. Our data further demonstrate that the majority of DNA evolution correlated with the genitalia has been shaped by negative selection or neutral evolution. This indicates that genital changes are predominantly facilitated by relaxation of constraints rather than positive selection, possibly due to the high pleiotropic nature of the morphogenetic genes.

## Introduction

How phenotypic novelties arise and the processes that govern their change are pillars of the evolutionary biology research. Studies on the *evo-devo* era have shown that determining how genes coordinate morphogenesis during development is a fundamental step to understand phenotypic evolution (Carroll 2008; Mallarino & Abzhanov 2012). A symbolic example is the segmental origins of insect wings, which has been debated for centuries with no consensus (Averof & Cohen 1995; Medved et al. 2015). The elegant *evo-devo* study by Linz & Tomoyasu (2018) with *Tribolium* beetles revealed a possible “dual origin” for these structures, suggesting wings are in part a modified leg and simultaneously a tergal expansion (Linz & Tomoyasu 2018). Undeniably, individual-level studies as such provide invaluable data to study phenotypic evolution. Nevertheless, single-species approaches are insufficient to explore the evolutionary processes that act after the emergence of novel traits. Our understanding of these processes are elusive, and we have poor knowledge on how the genetic-developmental architecture of morphological traits evolve themselves and how their evolution is mirrored in the phenotypic change across large time scales (Baker et al. 2021). There is an urgent need for more studies associating morphological, developmental and genetic data in comparative frameworks (Sanger & Rajakumar 2019).

An illustrative example of this gap is the study of genital evolutionary development in insects. There has been growing research scrutinizing the genes and pathways underlying genital development (e.g. Smith et al. 2020; Xu et al. 2019; Vincent et al. 2019). Notably, recent studies with *Drosophila* and *Carabus* beetles revealed sequence divergence of distinct genomic regions associated with the divergence of genital traits in recently separated species (Fujisawa et al. 2019; Hagen et al. 2021). These data may have started to illuminate the role of genital morphology and their underlying genetics in the speciation process. A direct prediction from these observations is that the outstanding rates of genital evolution in insects may have left genomic signatures over time. For example, we might expect that accelerated rates of genital diversification correlates with changes in the rates of molecular evolution or with the strength of molecular adaptation. One avenue to test this hypothesis is to estimate the degree of association between genitalia and molecular substitution rates across a phylogeny. The advantage of this approach in our understanding of the evolutionary development of genitalia would be two-fold. First, studying the genotype-phenotype association at the macro-evolutionary scale allows investigating the predominant mechanisms that have led to diversification (Lartillot & Poujol 2011; Jones et al. 2020). For instance, has rapid genital evolution been facilitated by positive selection acting in key developmental genes? Second, the comparative approach provides a way to evaluate the developmental and genetic bases of morphological evolution across an entire clade instead of a single species (Organ et al. 2015). In this study, we tackle this topic by reconstructing the evolution of male genital size across the order Hemiptera, spanning over 350 m.y. of evolution (Johnson et al. 2018). Following, we link the phenotype with the genotype by estimating the association between genital size and the rates of sequence evolution of several developmental genes.

Size is one of the most important attributes of genitalia in insects. A proper genital size must be guaranteed during development, since it allows the perfect fit between males and females and avoids mechanical mismatches in copula (Usami et al. 2006; Tanaka et al. 2018). As a result, genital size tends to develop distinctly from other traits, usually showing a negative allometric scaling - *i.e*. a relatively constant size despite variations in overall organism size (Lupše et al. 2016). There is solid evidence that genital size develops more independently through an insensitivity to systemic regulators of body size development, for example the insulin signaling cascade (Dreyer & Shingleton 2019; Tang et al. 2011; Emlen et al. 2012). While insulin insensitivity may help explain patterns of genital growth during ontogeny, which genes and pathways underlying this special mode of development of genital size are yet poorly understood (Terada et al. 2021). By extension, we know even less about the genetic-developmental mechanisms that may allow, facilitate or constrain genital size diversification in the evolutionary scale. For example, a handful of studies have shown that knocking down central developmental genes may result in smaller genitals, like genes involved in sexual determination, segmental organization and appendage patterning (Aspiras & Angelini 2011; Macagno & Moczek 2015). However, if those master regulatory genes or, alternatively, more specific downstream genes, are the ones whose structure are shaped by selection in the evolution of genital size, is largely unknown.

Here, we employ a comparative approach to investigate the evolution of genital size across 92 hemipteran species (Fig. 1), aiming at the evolution of its underlying genetic-developmental machinery. We use phylogenetic methods and publicly available genome-scale data to estimate the rates of evolution of 68 genital-development genes and correlate those rates with changes in genital size and development rate. We further evaluated the most likely selective regimes associated with these genes by reconstructing the rates of synonymous and non-synonymous substitutions. Our analyses revealed that genital size in hemipterans has evolved in dissociation with development rate but, in contrast with former observations, in close association with body size. We also found 18 genes whose substitution rates were significantly correlated with genital size. Lastly, we show that only a few sites in these genes have evolved under positive selective pressures, while the majority of sequence change has been driven by negative selection and neutral evolution. We discuss how our data fits or disagree with current theory of genital evolution, and how our analyses illuminate the mechanisms underlying the macro-evolution of genital size and its genetic architecture.

**Figure 1.**
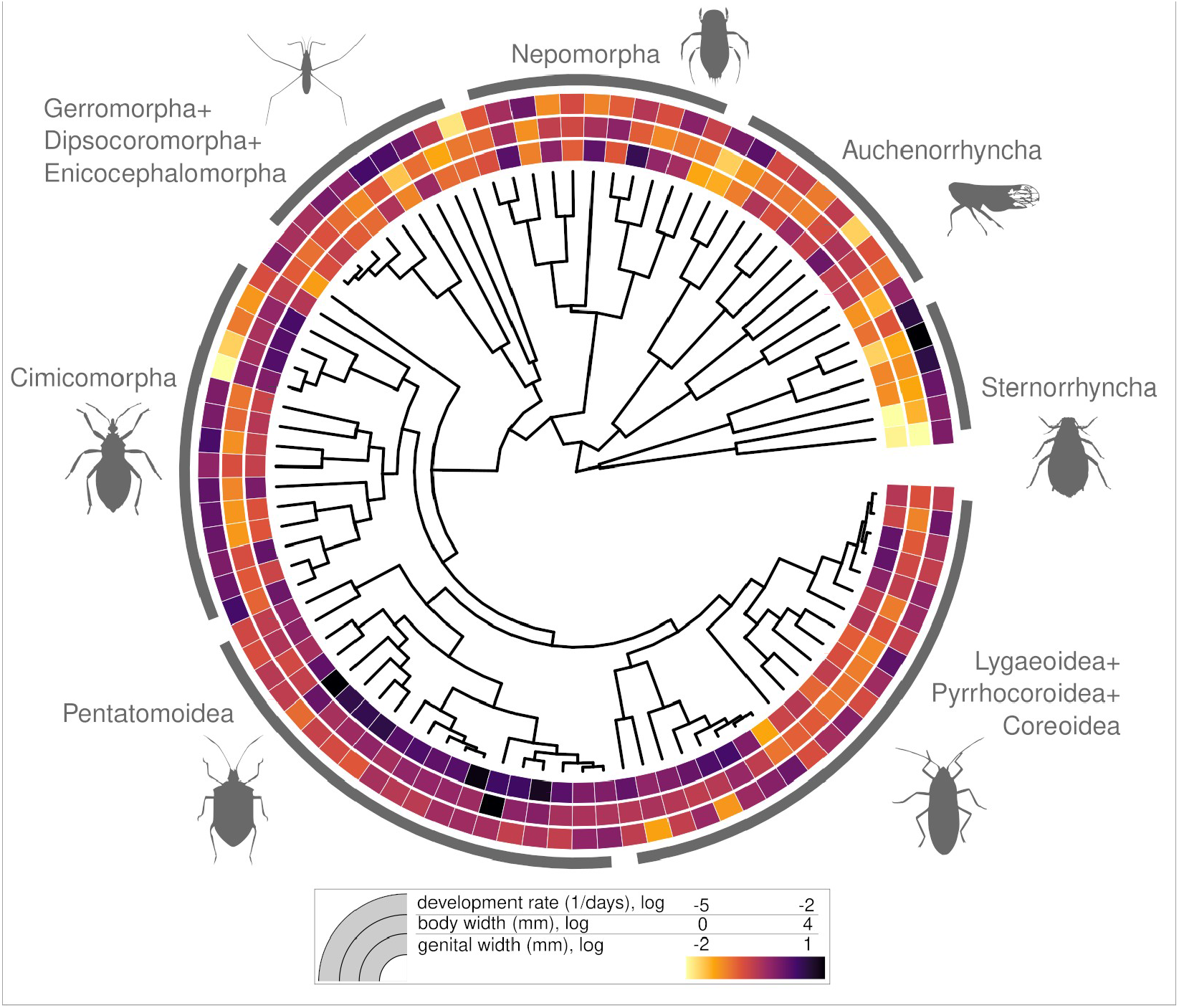
Phylogenetic diversity of genital size (inner circle), body size (middle circle), and development rate (outer circle) across the Hemiptera. There is a clear trend of larger species exhibiting larger genitals (e.g. Pentatomoidea and Sternorrhyncha), while development rate seems less correlated with both morphological traits.

## Material and Methods

### Phenotypic and developmental data

We used the maximum width of the external genitalia as a proxy for genital size and maximum abdominal width as a proxy of body size. The majority of measures were collected directly from scaled images published in the literature or in online databases, but a few species were measured from personal figures (Supplementary Data). All measures were taken in the software ImageJ.

We compiled information on the total post-embryonic development duration in number of days from the literature. Because different studies were conducted in different temperature conditions, we used the Campbell’s (1974) developmental model to standardize developmental duration and determine the developmental rate, defined as 1/(developmental time). The minimum temperature for successful development in Hemiptera usually varies between 10 and 16 degrees Celcius, thus, we used the median 13.5 for minimum developmental temperature in the model. We then calculated an adjusted developmental rate for a standard temperature of 25 °C. The vast majority of data was obtained from studies with controlled temperature (Supplementary Data). For the few studies conducted in the field or at room temperature, we used the mean month temperature where the study was conducted, taken either from the respective study (when reported) or from web databases (*climate-data.org* and *gspatial*).

### Genital-development genes and bioinformatics

We searched for genes associated with genital development in the platforms FlyBase, VectorBase, and The Gene Ontology Resource (GO:0035112 and GO:0048806). We identified a total of 68 genes for which we could obtain ortholog protein sequences to *Drosophila melanogaster* in the genomes of the bed bug (*Cimex lectularius*) and the kissing bug (*Rhodnius prolixus*). We used protein sequences from these two bugs as reference to search for the orthologs across all publicly available hemipteran transcriptomes.

We downloaded 76 hemipteran transcriptomes from the TSA database and assembled *de novo* eight transcriptomes with raw RNA-seq data obtained from the SRA database (Supplementary Data), totaling 92 species. We used Trinity v. 2.11.0 (Grabherr et al. 2011) for the transcriptome assembly, with minimum contig length of 199 bp and other parameters set to default. To search for the orthologs of the 68 genes in these transcriptomes, we used blastX in Diamond v. 0.9.21.122 (Buchfink et al. 2014), with a minimum e-value of 0.0001, 40% of query length coverage and 70% of identity. We further selected the best hit for each species and each gene filtering by bitscore with the python script “blastfilterer.py” (https://github.com/juancrescente/biopyutils/blob/master/blastFilter.py).

To detect protein-coding regions and conduct alignments we used AlignWise v. 0.38 (Evans & Loose 2015) with the muscle algorithm and TranslatorX v. 9.0 (Abascal et al. 2010) with amino acid-guided alignments. We did a manual curation of each alignment to remove misaligned and gap-rich sequences.

### Phylogeny

Since there is not a single phylogeny for all the taxa of interest, we used a super-tree approach in CLANN with the average consensus algorithm, which preserves branch lengths (Creevey & McInerney 2005). We inferred the supertree using eleven molecular phylogenies of hemipterans (Yan Hui Wang et al. 2017; Juan Wang et al. 2017; Forthman et al. 2019; Johnson et al. 2018; De Moya et al. 2019; Li et al. 2012; Cao et al. 2020; Liu et al. 2019; Zhang et al. 2016; Nováková et al. 2013), posteriorly replacing taxa with closely related species to match those with genital and transcriptomic data (Supplementary Data). We dated the phylogeny using the function “chronos” in the R package Ape v. 5.5 (Paradis et al. 2003), with a relaxed model and the same calibration points as in (Johnson et al. 2018). The resulting ultrametric tree was used for all comparative analyses.

### Comparative analyses

We started by evaluating the allometric pattern of genital size evolution. We first conducted a generalized least squares (GLS) analysis where log-transformed genital width was the response variable and the log of the major abdominal width was the predictor variable. Second, we also tested for the effect of development in the evolution of genital size by including in the first model the adjusted developmental rate as a second predictor variable. Third, we also included in these models a phylogenetic correlation matrix estimated under a brownian motion model to test the effect of phylogenetic non-independece (pGLS). We then compared these models using the Akaike information criterion (AIC).

To investigate the genomic signatures underlying genital size evolution, we modeled the evolution and coevolution of molecular substitution patterns and genital size in Coevol v. 1.5 (Lartillot & Poujol 2011). Briefly, the program models the evolution of synonymous (*dS*) and non-synonymous substitutions (*dN*) across a phylogeny, as well as the phenotypic data of interest, using a brownian motion model in a Markov Chain Monte Carlo (MCMC) approach. Here, we were particularly interested in the ratio *dN/dS (i.e*. omega, ω), which may inform about adaptation. The association between genital size and ω was estimated using a multivariate brownian motion model, where the strength of the association was assessed using the partial correlation coefficient R, and the significance was determined by a posterior probability. We interpret R > 0.5 as moderate correlations, R > 0.7 as strong correlations and R > 0.85 as very strong correlations. We adopted a significance threshold for the correlations of 0.95 of posterior probability. MCMC burn-in was set to 15%, while mixing and convergence were assessed using the module tracecomp within Coevol (Lartillot & Poujol 2011), where the analyses were stopped once a minimum effective sample size of 40 was achieved for the following parameters: log prior, log likelihood, tree length, *dN/dS* and the covariance matrix (Lartillot & Poujol 2011).

We next determined the most likely selection regime acting on the genes that showed significant association with genital size, *i.e*., negative selection, neutral evolution, or positive selection. For this, we fit each alignment to a site model (M2a) in CodeML from PAML v. 4.9 (Yang 2007), which estimates omega independently for each site (*i.e*. codon). We used the same tree from previous analysis and the Bayes Empirical Bayes criterion to determine the selection regime.

## Results

Our literature compilation revealed great variation in genital and body size as well as in development rate in Hemiptera (Fig. 1). All traits were highly heterogeneous across the phylogeny. For example, genital size exhibited the largest values and were relatively constant in pentatomoids, while extremely variable in nepomorphans and lygaeoids + allies. There is a clear visual trend of larger species exhibiting larger genitals (Fig. 1), which is mirrored by our models. The simplest GLS model, where genital size is explained exclusively by body size, revealed a significant correlation (R^2^ = 0.64, p = 2e^−16^, Fig. 2). This “pure allometric model” was the best model (Table 1), meaning that including the development rate and the phylogeny as explanatory variables, either in combination or separated, always decreased model fitness (Table 1). These results indicate that genital size evolution is weakly influenced by development rate and has low phylogenetic signal across the Hemiptera order (Figs. 1–2, Table 1).

**Figure 2.**
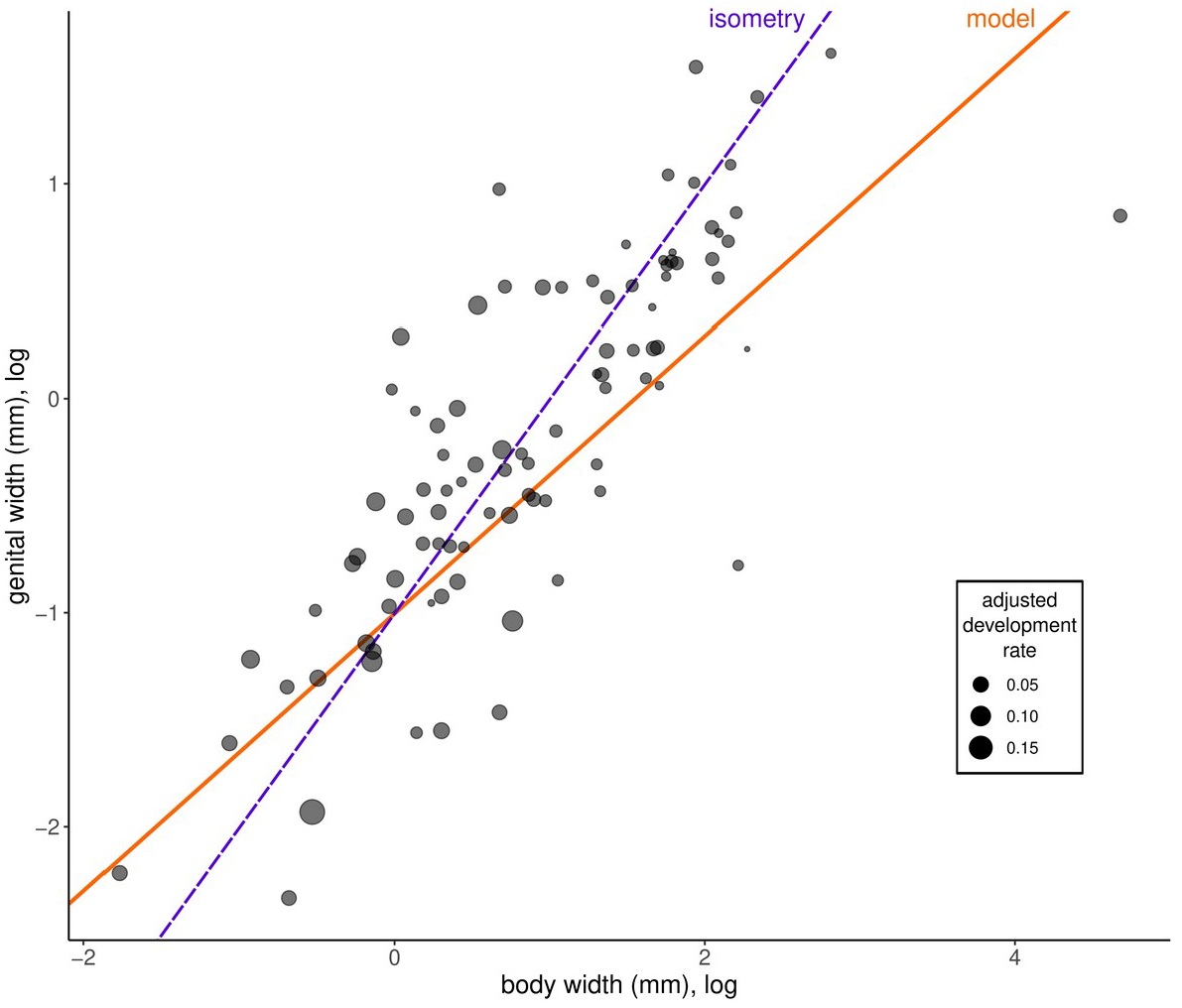
Relationship between body and genital size (given in mm, log transformed), where each point is a hemipteran species. Best fit line (orange) is from the best model, the phylogenetic generalized least squares analyses including the development rate (point size) as a predictor variable. Theoretical isometric relationship is expressed in purple.

**Table 1.**
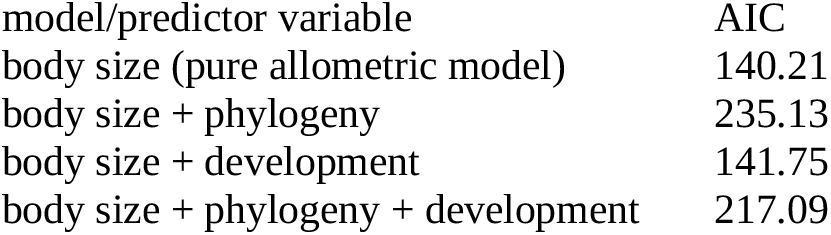
Results of generalized least squares modeling.

| model/predictor variable | AIC |
| --- | --- |
| body size (pure allometric model) | 140.21 |
| body size + phylogeny | 235.13 |
| body size + development | 141.75 |
| body size + phylogeny + development | 217.09 |

Since body size had a strong effect on genital size, we used the ratio between these two variables to test for the genotype-phenotype correlations. This approach allowed us to discount body size, modeling only body-size-free genital variation instead. For the 68 genes tested, 18 genes showed a significant correlation between omega (ω) and corrected genital size (posterior probability > 0.95) (Fig. 3). Out of these 18 genes, three exhibited very strong positive correlations (*cabeza, escargot* and *homothorax*), and one exhibited a very strong negative correlation (*costa*) (Fig. 3). Seven genes showed strong correlations, while nine genes had moderate correlations (Fig. 3). A full table of the Coevol results is provided as Supplementary data.

**Figure 3.**
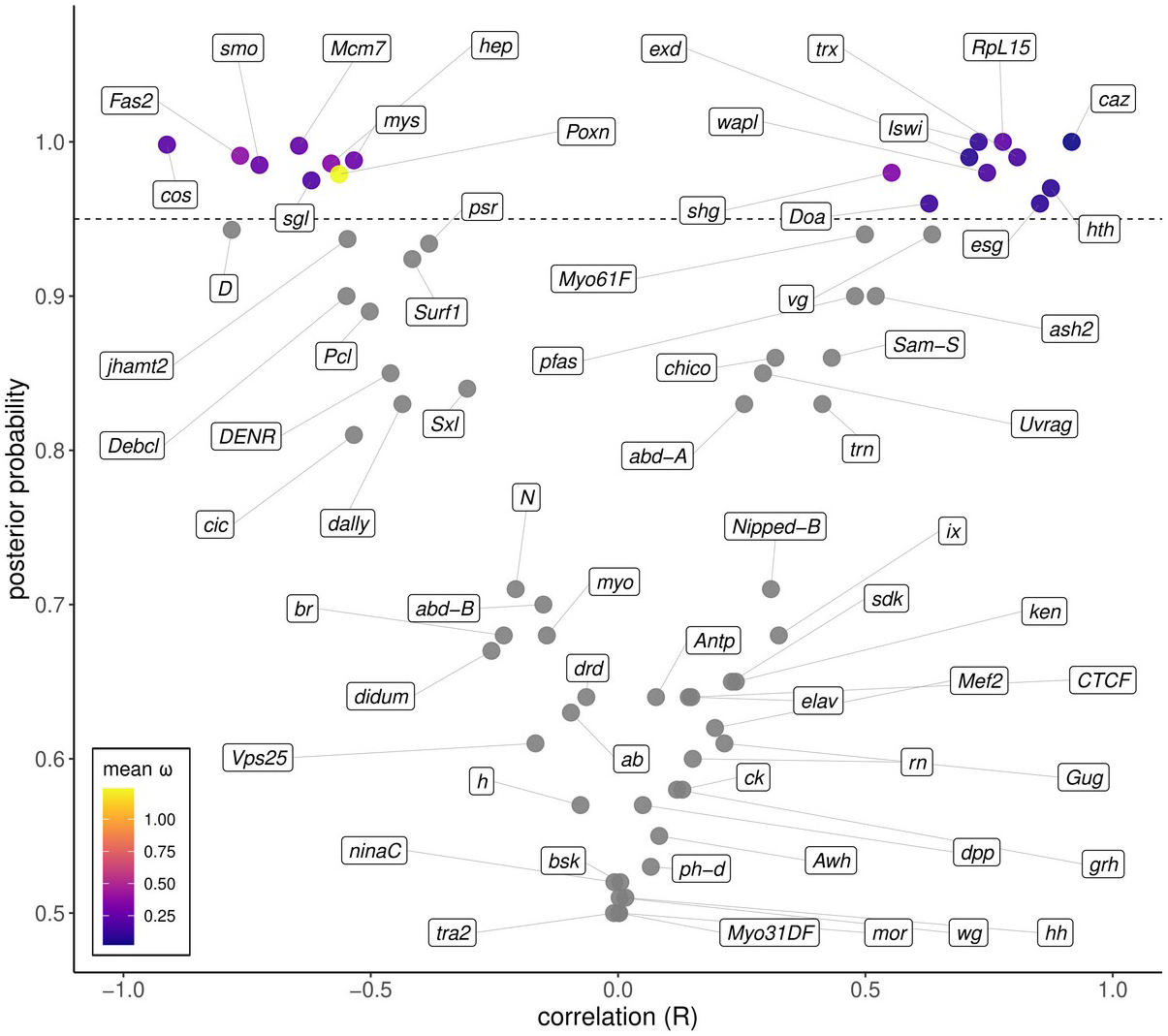
Results of phenotype-genotype association (body-size-corrected genital width versus omega (ω)) estimated in Coevol, showing the degree of correlation versus the posterior probability (y axis). Each point represents a gene, and the dashed line is the posterior probability cut-off of 0.95. Omega values for each gene with significant correlation with the phenotype were taken from the site-wise model from CodeML, thus representing the mean omega of all codons.

For the 18 genes with significant association with genital size, the analyses in CodeML revealed that nine genes had only negatively selected or neutral sites (Fig. 4). The majority of sites of the other nine genes showed a prevalence of negative selection and neutral evolution, with only a few sites positively selected (Fig. 4). For these genes, the number of positively selected sites varied from two (*costa, smoothened* and *sugarless*) to nine (Pox neuro and shotgun). Omega values for the positively selected sites ranged between 1.37 (costa) and 8.09 (Pox neuro).

**Figure 4.**
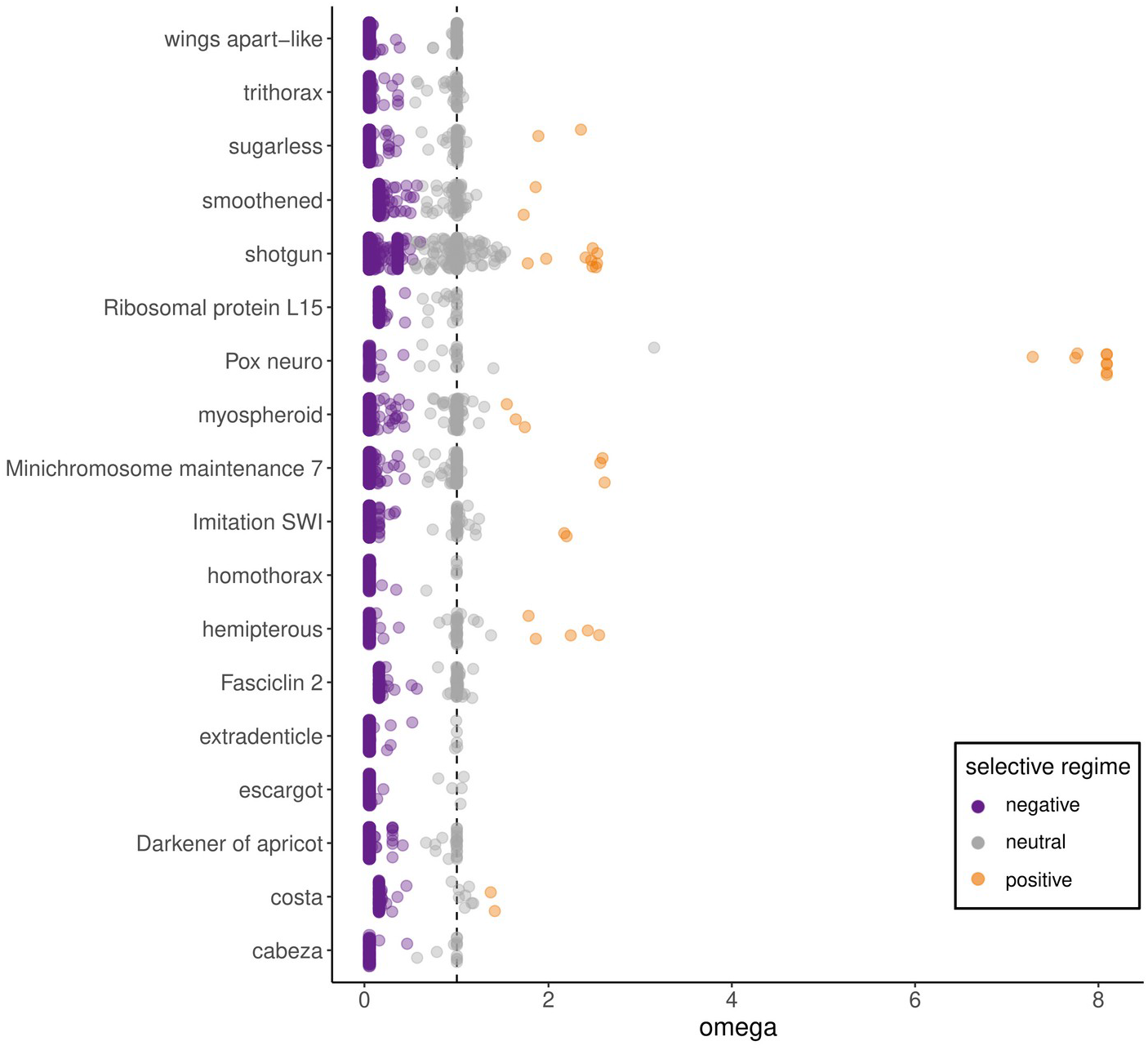
Selective regime estimated in CodeML for each codon (colored points) of all genes that showed significant association with genital size (detected in Coevol). The vertical dashed line approximates the positive selection zone threshold, where *dN/dS* are equal (ω = 1).

## Discussion

In this study, we conducted the largest and most comprehensive analysis of genital size evolution, with samples representing an entire insect order, the Hemiptera. Below, we first discuss the morphological patterns we revealed framed into the phylogenetic perspective, considering how our data fits the predictions of the theory of genital evolution. Second, we discuss our findings of the adaptive molecular evolution analyses, focusing on differences observed in substitution rates for different genes, as well as their implications for genital *evo-devo*. Lastly, we provide a perspective on how our approach can be expanded to strengthen our understanding of these genotype-phenotype correlations.

### Constraints of body size and development rate in genital evolution

We showed that there is great variation in genital size across the Hemiptera, and the magnitude of among-species variation may vary across clades. This observation is not surprising and it is consistent with recent studies that revealed heterogeneous patterns of genital change in beetles (Rudoy & Ribera 2016; Genevcius et al. 2020). Nevertheless, we still observed a clear allometric signal, meaning that the evolution of genital size is constrained by body size to some extent. In principle, this result may seem contradicting with the theory of genital allometry, which postulates that genital traits develop more independently of body size than other traits. In simple terms, the rationale is that genital traits should be less sensitive to size variations to avoid mismatches between males and females (House & Simmons 2007). We emphasize, however, that allometry is a phenomenon that takes place at the populational/individual level, where organ growth may be more or less restricted to body growth (Mirth et al. 2016). Most of the studies that found negative allometry for genital traits have compared body and genital sizes either among individuals of a single species or among ontogenetic stages (Orbach et al. 2018; De-Lima et al. 2019). Predicting and testing how allometric scaling evolves itself, or, in other words, extending our knowledge of the ontogenetic allometry to evolutionary time, is much less straightforward (Pélabon et al. 2014). Accordingly, the few studies evaluating genital allometry in broader scales have found more complex scenarios. For example, Galicia-Mendoza et al. (2021) found distinct allometric patterns for the genitalia of aggressive and non-aggressive damselfly species. In spiders, it has been shown that non-intromittent genitalia are evolutionarily correlated with body size, while intromittent genitals are not (Lupše et al. 2016). In Hemiptera, the within-species variation of body size is, at most, modest. In contrast, among-species variation of body size is enormous within the order (see Fig. 1). Taking species of our analyses as examples, the aphids (Sternorryncha) are usually 2-4 mm in body length, while some pentatomoids may reach over 5 cm. Therefore, even if the genitalia in most of our studied species were relatively insensitive to body size during development, it is certain that the enormous variation of body sizes across the the order has been a constraint for the evolvability of genital size.

Genital size was also found to be weakly influenced by development rate. This result was largely expected due to functional and ontogenetic characteristics of the structure. Since immatures do not copulate, genitalia are the latest structures to develop during ontogeny, together with wings. External genitals in hemipterans develop mostly during the fifth and last nymphal instar, and are only completely formed in adults (Singh 1971). Therefore, unlike other structures that are functionally important since early development such as the mouth parts, genitalia are completely absent in the stages where we observe massive body size increase, i.e. during the first four molts.

### Molecular signatures of genital macro-evolution

We investigated the evolutionary patterns of 68 genital-related genes across 92 hemipteran species and demonstrated that, for 18 genes, the ratio of non-synonymous and synonymous mutations (dN/dS or ω) correlated with genital size. None of these genes were master regulators of signaling pathways, like the hox genes that determine segmental identity of posterior body (e.g. Abd-a, Abd-b, Antp), nor sex determining genes (e.g. Ix and Sxl). As expected, such genes had low correlations with the phenotype, probably because their coding regions are highly conserved (Krumlauf 2018), in contrast with the rapidly evolving genitalia (Genevcius et al. 2017). For these genes, changes in sequence composition would likely yield drastic phenotypic outcomes and be highly deleterious during development (Chen et al. 2022; Zhang et al. 2021). Another group of genes that showed overall low correlation with genital size were the appendage-patterning genes like decapentaplegic, hedhehog and wingless. Genitalia are commonly thought to be serially homologous to other appendages like legs and antennae due to demonstrated role of these appendage patterning genes in genital growth in holometabolous (Macagno & Moczek 2015; Estrada & Sánchez-Herrero 2001). In contrast, our results demonstrate that there are no signatures of an adaptive process acting on these genes concerted with genital size. Our data seems more in line with either of the two hypothesis that 1) the male genitalia in Hemiptera may be a highly derived appendage primordia which lost appendage patterning functions, or 2) genitalia are modified abdominal segments with absolutely no appendage homology (Aspiras et al. 2011). Note that it is still possible that appendage patterning genes may correlate with genitalia in the level of expression (see discussion below).

We found significant phenotype-genotype correlations for genes previously known to be associated with genital size. For example, knock-down in Homothorax has caused a reduction in copulatory organs in the milkweed bug (Aspiras et al. 2011); in *Drosophila*, Mcm7 knock-down has been associated with changes in clasper size (Tanaka et al. 2015), while Costal2 mutants showed increased claspers and reduced hypandrium (Sánchez et al. 1997). To some extent, the concordance between these ontogeny-level studies and our results at the macro-evolutionary scale provides validation to our approach. This is especially important given the paucity of studies linking evolutionary processes measured at the macro scale and direct experimental observations of gene function (Moury & Simon 2011). Interestingly, Mcm7 has been shown important in the development of genital traits with very recent divergence in *Drosophila* (Tanaka et al. 2015; Hagen et al. 2021). Taken together, these observations and our own results may indicate this gene as a candidate involved not only in morphogenesis but also in reproductive isolation and lineage diversification.

Another interesting aspect related to these “genital size genes” is that they are involved in completely different developmental pathways with distinct molecular and biological functions. For example, Homothorax is a TALE homeobox transcription factor that forms the *homothorax-extradencticle* route, promoting cell division and cuticle formation (Rieckhof et al. 1997). In contrast, Costal2 participates in the Hedgehog pathway and it is directly involved in growth control (Sánchez et al. 1997). In addition, we also observed significant correlations with genes related to other features of genitalia apart from size: bristle patterning, sclerotization, pigmentation, and the formation of several accessory structures (Table 2). Taken together, these results highlight that genital size is not a “simple trait” whose evolution has been merely facilitated or accompanied by changes in pathways that regulate overall body size. Rather, our results suggest that genital size evolution is only possible with accumulated changes associated with diverse morphogenetic processes, for instance, the formation of cuticle composition, sensory apparatus, and muscle development. Nevertheless, understanding why the evolution of genital size may be linked with the evolution of these other features is not straightforward. A plausible explanation is that the adaptive value of bigger genitals may be indirect. Since we measured the size of the genital capsule, which revolves the remaining genital structures, larger genitals may be associated with increased complexity (Song 2009), by allowing the accommodation of increased number of structures common in insect genitalia, such as hooks, setae, folds, etc. The development and evolution of these structures in more complex genitalia would predict selection in distinct molecular and biological functions, as suggested by our data.

**Table 2.**
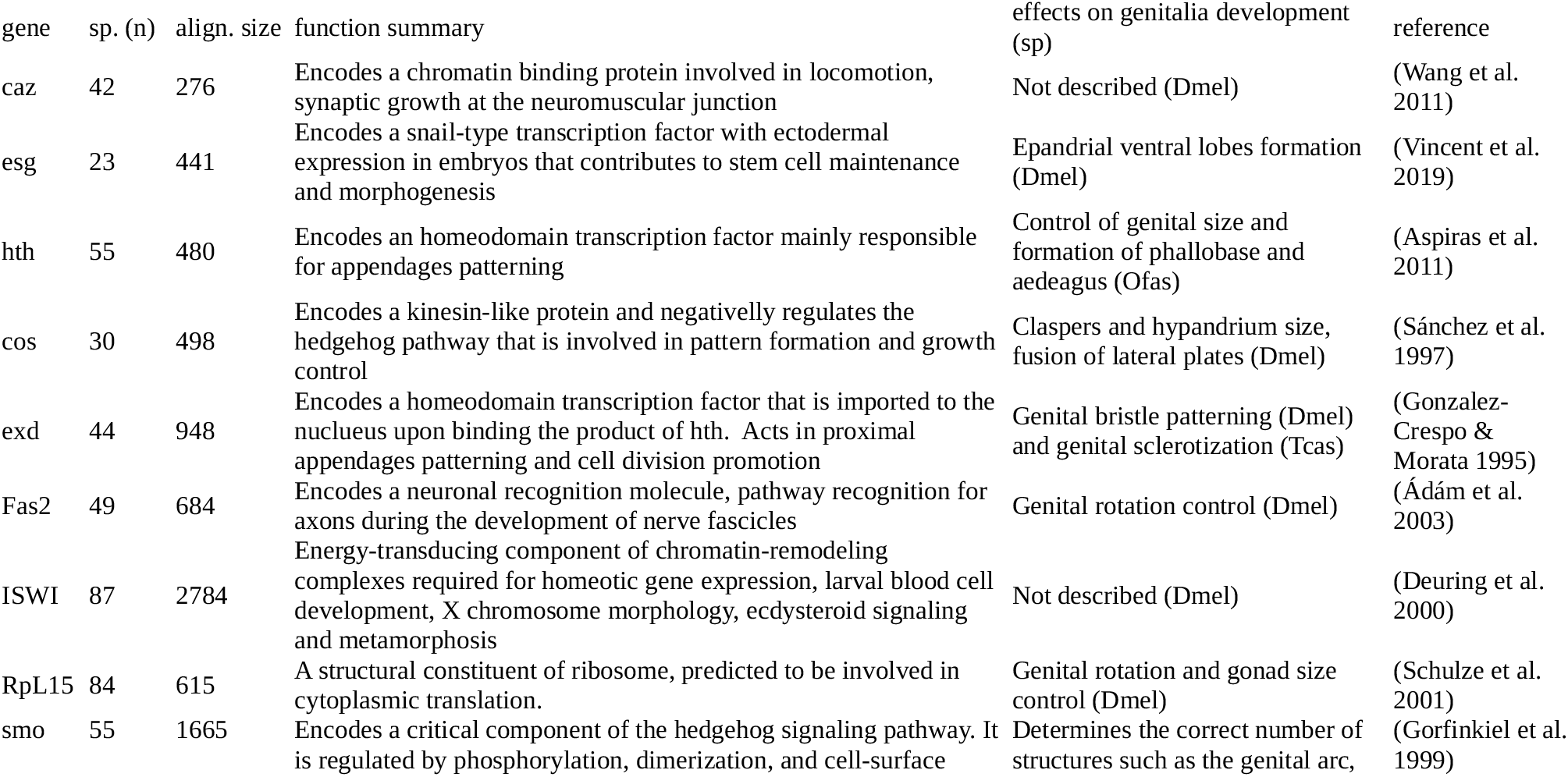
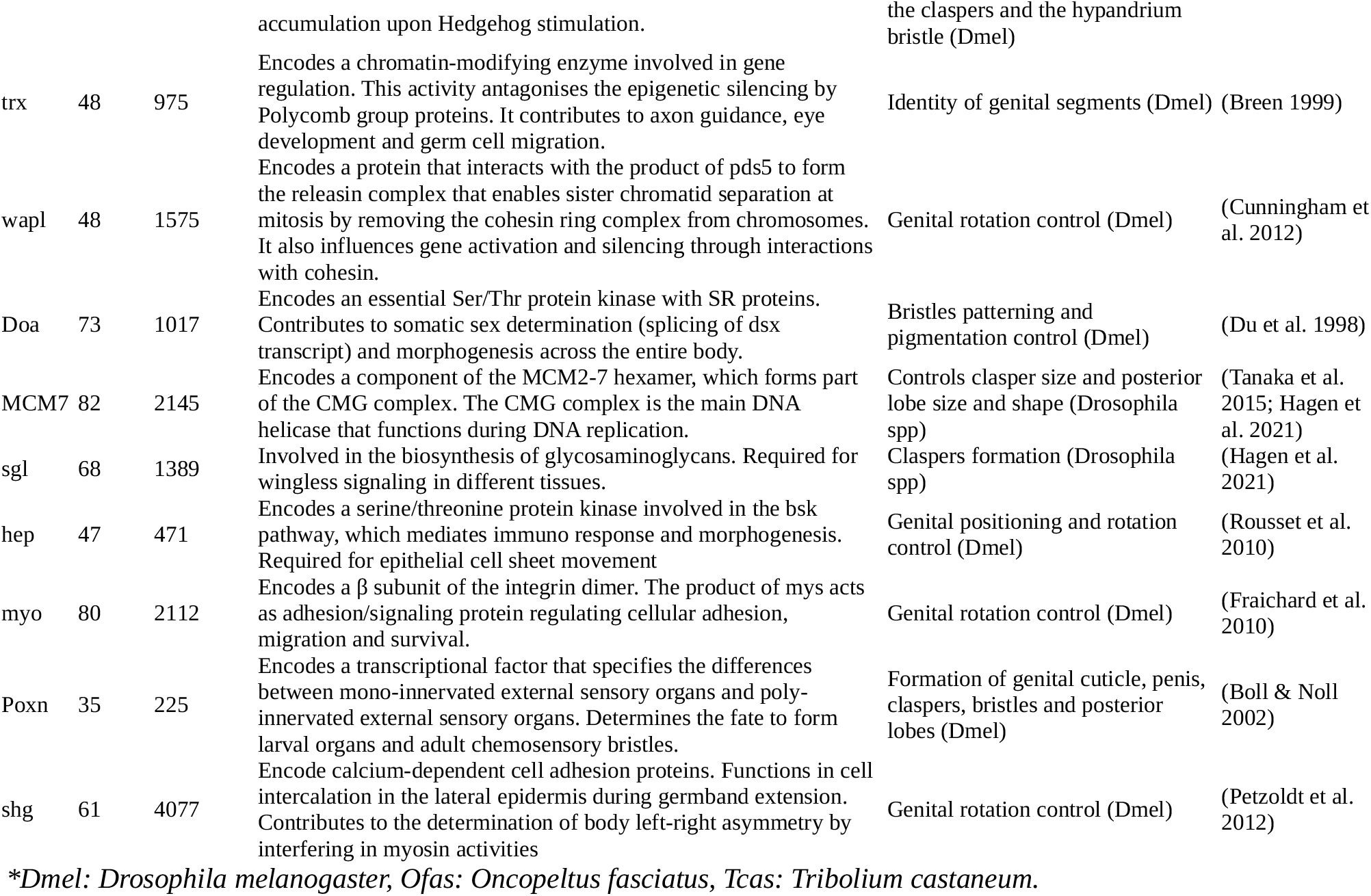
The 18 genes whose *dN/dS* were found to have a significant correlation with genital size. Because we did not detect orthologs for all species for all genes, each analysis had a different number of species (“sp” column). Final alignment size (align. size) used in Coevol analyses is denoted in nucleotides. Gene function and its phenotypic effects are described with its respective reference.

Although we observed very strong correlations between genital size and ω for some genes, we found that mean cross-species ω values were overall low (Fig. 3). This is consistent with a scenario where the majority of changes in genital size have been possible due to a relaxation of purifying selection in developmental genes rather than by persistent positive selection. In fact, this scenario is probably more often the rule then the exception due to the highly pleiotropic nature of the morphogenetic genes (e.g. Partha et al. 2017; Womack et al. 2018). From our dataset, illustrative examples are the transcription factors escargot, cabeza, and homothorax. Among different structures, these genes also mediate wing morphogenesis (Casares & Mann 2000; Fuse et al. 1996), which is a phenotype that seems highly conserved across Hemiptera. Still, for half of the genes that correlated with genital size, we observed at least a few sites under positive selection (Fig. 4). This highlights the possibility of heterogeneous evolutionary mechanisms acting in different protein regions, and such differentially affected regions may be interesting targets for future functional studies. It is also important to emphasize that synonymous mutations are not necessarily dissociated with adaptive evolution as they may be advantageous in some contexts like changing splicing patterns and the speed of RNA translation (Chu & Wei 2019). This topic is another interesting avenue for future studies.

One last key question that emerges from our results is why some genes exhibit positive correlations with the phenotype while others exhibit negative correlations (see Fig. 3). While this question can only be precisely answered with deeper understanding of gene functions coupled with the underlying phenotypic selective mechanism, we hypothesize that genes with negative correlations may represent “moderator genes” (Baker et al. 2021). In this scenario, mutations in protein-coding regions would be thought to consistently decrease phenotypic change. This may be particularly the case of genes that act as negative regulators of developmental pathways such as Costa and Fasciclin 2, which are involved in transcription factor sequestering of the Hh pathway and the inhibition of epidermal growth, respectively (Mao & Freeman 2009; Li et al. 2016). In addition, genes with marked pleiotropic and epistatic behaviors may also act as moderators due to their global effects on developmental pathways, such as smoothened and sugarless.

### Future directions

Our approach has been shown interesting to reveal the genes and potential loci within these genes involved with the evolution of genital size. We show that, out of 69 genes analyzed, 18 genes had positive signatures of genotype-phenotype correlation. We emphasize, however, that associating phenotypes with mutation rates in coding regions, although powerful, only represents partial picture of the adaptive mechanisms underlying (macro)evolution. We thus highlight four directions for future research that may provide further advance on these mechanisms.

The first direction draws from the observation that we cannot completely discard the relevance of the genes without significant correlation because, apart from sequence structure, other features may evolve themselves: expression levels (Romero et al. 2012), expression timing (Keyte & Smith 2014), regulatory sequence composition (Hoekstra & Coyne 2007), structure of gene regulatory networks (Smith et al. 2018), among others. Measuring (co)expression levels of candidate genes in species with different relative genital sizes, and modeling these features across the phylogeny, represent an interesting future avenue. Secondly, it is important to note that our prior hypotheses of genotype-phenotype associations derive mostly from studies on *Drosophila melanogaster*. There are certainly other important genes and pathways that can be discovered by conducting genomic functional studies in non-traditional model insects, such as the Hemiptera. Third, an important characteristic of COEVOL and similar analyses is that they estimate a single omega value for the entire gene. This is not a problem *per se*, but it may exclude genes for which single sites may coevolve with the phenotype, thus, only genes with a global correlation will be captured. To our knowledge, no study has yet used the COEVOL methods to estimate covariances between phenotypes and isolate codons. Even though there will be computational and theoretical challenges, this is another promising approach. Lastly, one crucial aspect of genotype-phenotype modeling is the assumption that this correlation is unidirectional (Smith et al. 2015), *e.g*. increases in omega leading to increases in trait size. However, it is possible that changes in omega rates may implicate heterogeneous phenotypic changes across different clades, making the detection of the correlation more difficult. A very recent method has been developed to account for this issue (Baker et al. 2021), and it may applied for future studies on genital evolution as well.

## Acknowledgements

We are grateful to Dr. David A. Rider for providing many of the articles used to gather morphological and developmental data, and Gisele A. Cardoso for valuable comments on the manuscript and coding tips. We also thank FAPESP for a postdoctoral fellowship to BCG (process n. 18/18184-4).

## Author Contributions

BCG and TTT designed the research. BCG and DCC collected the data. BCG analyzed the data. BCG, DCC and TTT interpreted the results and wrote the paper.

